# *The human GRK4γ 65L variant* causes salt-sensitive hypertension by increasing renal SLC4A5 expression through the HDAC1 pathway

**DOI:** 10.64898/2026.03.09.710443

**Authors:** Santiago Cuevas, Selim Rozyyev, Hewang Lee, Celia Arias-Sánchez, Daniel Yaqub, Jun Feranil, Prasad Konkalmatt, Raisha Campisi, Jacob Polzin, Laureano D. Assico, Ines Armando, Pedro A. Jose

**Author notes:** **Corresponding authors:** Santiago Cuevas PhD, Principal Investigator Miguel Servet, Physiopathology of the inflammation and oxidative stress lab | Molecular Inflammation Group, **BioMedical Research Institute of Murcia (IMIB-FFIS),** Laib Building, lab 4.30, office 5.2. Ctra. Buenavista s/n,E-30120 Murcia (Spain) www.inflammation.imib.es|. **Translational Statement:** The human GRK4γ 65L variant promotes salt-sensitive hypertension by increasing renal SLC4A5 and AT1R expression through HDAC1 inhibition. These molecular alterations enhance sodium reabsorption in the kidney, impairing the normal natriuretic response to high salt intake. Our findings in multiple mouse models and human renal cells reveal a conserved pathogenic mechanism linking GRK4 and SLC4A5 variants to dysregulated sodium handling. This pathway identifies HDAC1-dependent epigenetic modulation as a potential therapeutic target in salt-sensitive hypertension. These insights may guide personalized treatment strategies for patients carrying GRK4 or SLC4A5 risk alleles.

## Abstract

Salt-sensitive hypertension, a condition in which the blood pressure (BP) increases with an increase in salt intake, is influenced by behavioral, genetic, and environmental factors. Salt sensitivity is associated with variants of the G protein-coupled receptor kinase 4γ (GRK4γ) and the renal sodium bicarbonate cotransporter 2 (NBCe2), encoded by the *solute carrier family 4 member 5* (*SLC4A5*). The R>65L variant (*rs2960306*) of human GRK4 (*hGRK4γ 65L*) contributes to salt sensitivity through a signaling pathway and gene-gene interaction with *SLC4A5*.

Global expression of *GRK4γ 65L* in transgenic mice results in salt-sensitive hypertension, due in part to an increase in endogenous GRK4 and angiotensin type 1 receptor (AT_1_R) expression. Grk4 knockout (*Grk4^-/-^*) mice have decreased blood pressure and are salt-resistant. The expression of *hGRK4γ 65L* only in the kidney of *Grk4^-/-^* mice increases BP in response to a high salt diet. The renal expression of *SLC4A5* is increased in *hGRK4γ 65L* transgenic mice, relative to mice expressing wild-type (WT) human *GRK4* (*hGRK4 65L*), without endogenous mGrk4. Human renal proximal tubule cells (hRPTCs) endogenously expressing *GRK4 WT* and *SLC4A5 WT*, *SLC4A5* variants, *GRK4 65L,* and both *GRK4 65L* and *SLC4A5* variants were studied. SLC4A5 expression is increased in hRPTCs expressing *GRK4 65L* and in cells expressing both *GRK4 65L* and *SLC4A5* variants compared with *GRK4 WT* and *SLC4A5 WT.* Luminal and basolateral sodium transport in hRPTCs is increased in the presence of both *hGRK4 65L* and *SLC4A5* variants. GRK4 interacts with nuclear histone deacetylases (HDACs). Mice expressing *hGRK4 65L* only in the kidney have decreased expression but increased phosphorylation of HDAC1. HDAC1 expression is decreased and HDAC1 but not HDAC2 phosphorylation is increased in hRPTCs expressing both *hGRK4 65L* and *SLC4A5* variants. The presence of *hGRK4γ 65L* decreased HDAC1 expression but increased AT_1_R expression in the kidneys of mice on high salt diet. Our results show that GRK4γ 65L causes salt-sensitive hypertension by increasing renal SLC4A5 and AT_1_R expressions by inhibiting the HDAC1 pathway.

## Introduction

The long-term regulation of blood pressure (BP) rests on renal and non-renal mechanisms (1). The impaired renal Na^+^ handling in essential hypertension and salt sensitivity may be caused by aberrant counter-regulatory natriuretic/antinatriuretic pathways (1–3). The sympathetic nervous and renin-angiotensin-aldosterone systems are examples of antinatriuretic pathways (1, 3, 4). An important counter-regulatory natriuretic pathway is afforded by the renal autocrine/paracrine dopaminergic system, aberrations of which are involved in the pathogenesis of hypertension (5). The increase in BP with an increase in Na^+^ intake occurs in normotensive as well as hypertensive humans and predicts increased number of cardiovascular events and mortality, irrespective of basal BP levels (6, 7). Salt sensitivity is estimated to be present in 51% of the hypertensive and 26% of the normotensive populations (8). The mechanisms underlying salt sensitivity are not well understood.

G protein-coupled receptors (GPCRs) are the largest group of membrane receptors in eukaryotes, expressed in diverse cell types (9), and involved in numerous physiological processes, such as regulation of renal function and BP (10). The activity of GPCRs is regulated by G protein-coupled receptors kinases (GRKs) and despite the approximately 800 genes encoding GPCRs in the human genome, only seven GRKs (GRK1–7) have been identified (11, 12). The seven mammalian GRKs are classified into three sub-groups: the GRK1-like subfamily, the GRK2-like subfamily, and the GRK4-like subfamily, which includes GRK4, GRK5, and GRK6 (11, 12). GRK1, GRK4, and GRK7 have very limited distribution. GRK1 and GRK7 are found almost exclusively in the retina, and GRK4 is present in a few organs or tissues (11–13). Renal GRK4 is involved in the regulation of various cellular processes by phosphorylating GPCRs, such as dopamine receptors which are particularly important in the regulation of sodium balance and BP (14). Several variants of the GRK4 gene have been identified in humans. The main variants are due to alternative splicing, resulting in four splice variants: GRK4α, GRK4β, GRK4γ, and GRK4δ which differ in their amino acid sequences and functional properties (15).

Abnormalities in GRK4 function can lead to disorders such as hypertension due to its role in modulating renal function and sodium reabsorption (14); variants in the promoter region of *GRK4* can influence its expression (16) and the salt sensitivity of C57Bl/6J mice is related to increased renal expression of GRK4 (17). Renal-selective silencing of *Grk4* decreases the BP in the SHR rats (18); GRK4 interacts not only with D_1_R but also AT_1_R and ETBR in the regulation of BP (19–22). *GRK4* human variants (R65L, A142V, and A486V) have been associated with hypertension and/or salt sensitivity in several ethnic groups (23, 24). We have reported that intronic variants (intron 22-23 [**rs7571842**] and intron 25-26 [**rs1017783**]) of *SLC4A5* and *GRK4* (*GRK4 R65L* [**rs2960306]**) are associated with salt sensitivity in two Euro-American populations (23), thus, the treatment of hypertension can be gleaned from the pharmacogenomics of GRK4.

The *A142V* and *A486V* variants of *GRK4* are single nucleotide polymorphisms (SNPs) that have been reported by our group to be linked to the regulation of BP (8, 14, 19, 25). The *A142V* variant involves the substitution of alanine with valine at position 142, while the *A486V* variant involves a similar substitution at position 486. These mutations increase the kinase activity of GRK4, and we have reported that A142V increases the expression of AT_1_R and impairs the function of the renal D_1_R and D_3_R; its presence increases BP in mice fed a normal salt diet (25, ^26^). By contrast, the expression of A486V GRK4 variant in mice impairs renal dopamine receptor function and causes renal oxidative stress leading to impaired sodium excretion and elevated BP only in mice fed a high salt diet (10). However, the role of the R65L GRK4 variant in the pathogenesis of hypertension has not been reported.

The lack of powerful genetic associations in essential hypertension, especially salt-sensitive hypertension could be taken to indicate the importance of gene modification, such as that in epigenetics, especially resulting from environmental influence (27–30). Salt intake can increase oxidative stress and oxidative stress can influence epigenetics (e.g., HDAC activity) (27, 28). D_1_R, D_2_R, and D_5_R are important in the maintenance of normal redox balance and aberrant function of dopamine receptors can result in renal oxidative stress and hypertension (31–34).

All the above evidence points to the importance of GRK4 on BP regulation, oxidative stress, and HDAC1 function, and its capacity to regulate the expression and activity of key genes associated with hypertension such as SLC4A5, AT_1_R, and dopamine receptors. The aim of this study is to determine the consequences of the presence of human GRK4 65L on BP regulation and its interaction with other genes associated with hypertension

## 2. Material and Methods

### Animal Models

#### Transgenic hGRK4γ WT and hGRK4γ 65L

Transgenic mice expressing hGRK4γ^WT^ and hGRK4γ^65L^ were generated at the University of Michigan Transgenic Core Facility, using a protocol similar to that used to generate hGRK4γ 142V and hGRK4γ 486V transgenic mice (10, 25, 26, 35). The mice were crossbred with SJL and C67Bl6/J mice. The genetic background of mice used in the current study is 25% SJL and 75% C67Bl6/J (27) or 100% C67Bl6/J. Age-and sex-matched 3-to 8-month-old mice were studied.

#### Generation of AAV GRK4γ WT and hGRK4γ 65L mice

AAV vectors carrying hGRK4γ WT (AAV GRK4γ WT) or the mutant hGRK4γ 65L (AAV GRK4γ 65L) were constructed with the expression driven by a cytomegalovirus (CMV) promoter using the plasmid pAV-FH (Vigene Bioscience Inc., Rockville, MD, USA), as previously described (49, 50). These AAV-9 vectors were designed specifically to deliver target genes in transgenic mice, with one vector encoding human GRK4γ WT and the other encoding the human GRK4γ 65L variant.

We used six-month-old *Grk4^-/-^* mice (generous gift from Dr. Richard T. Premont, Duke University) to express selectively in the kidney hGRK4γ WT and the hGRK4γ 65L variant in a mouse lacking the mouse *Grk4*. We also used AAV carrying a control vector (CAAV) or human wild-type *SLC4A5* gene. The CAAV contains an EGFP cDNA in reverse orientation under the control of the CMV promoter and does not code for any protein. The mice underwent bilateral retrograde ureteral infusion of approximately 100 μl of AAV vector solution (1 × 10^11^ viral genome particles) to ensure kidney-targeted delivery (62). The BPs were measured prior to AAV infusion and 14 after AAV treatment after which time the organs were harvested and properly stored at 4°C for future analysis. The mice were euthanized with an overdose of pentobarbital (100 mg/kg).

#### Generation of hGRK4γ WT and hGRK4γ 65L mice

*hGRK4γ WT* and *hGRK4γ 65L* knock-in mice were generated at ViewSolid Biotech Inc. Knock-in of human genes into the mouse *Grk4* locus was performed through CRISP/Cas9-mediated homology-directed repair in C57Bl/6J mice. The gRNA Grk4-g4 was designed to target the start codon region of mouse *Grk4* to guide Cas9 to introduce a double strand-break that is then homology-directed repaired using donor template DNA containing either the *hGRK4γ WT* or *hGRK4γ 65L* into the mouse *Grk4* locus. The human genes are expressed under the mouse endogenous *Grk4* transcriptional control, in mice lacking the mouse *Grk4*.

#### Animal Procedures

Mice were fed either a normal salt diet (NS), containing 0.8–0.9% NaCl, or a high salt diet (HS), containing 6% NaCl, for a duration of 3 weeks. The mice had free access to food and water and housed in a temperature-controlled facility with a 12-hour light/dark cycle, and ambient temperature kept between 20°C and 26°C.

On the 21st day of the respective diets, urine samples were collected over a 24-hour period in mice kept in metabolic cages to measure sodium and creatinine levels. Following sample collection, the mice were anesthetized with an intraperitoneal injection of ketamine/xylazine (0.1 mg/100 g).

### Blood pressure measurement

The mice were anesthetized with an intraperitoneal injection of pentobarbital sodium (50 mg/kg). CardioM MaxII system (Columbus Instruments) was used to record BP from a catheter inserted into the femoral artery that was advanced into the abdominal aorta.

### Telemetry

Mice were anesthetized with an intraperitoneal injection of pentobarbital sodium (50 mg/kg). A telemetry transmitter (cat# TA11PA-C10, Data Sciences International) was magnetically activated 24 hours prior to implantation and soaked in warm saline for 10 minutes before surgery. The catheter end of the transmitter was inserted into the temporarily occluded left carotid artery, while the main body of the device was secured in a subcutaneous pocket created through blunt dissection of the right flank. The incision was closed using 4-0 Ethilon® sutures, and post-operative analgesia was provided with buprenorphine (0.05–0.10 mg/kg, intraperitoneally) (17).

Following surgery, the mice were monitored closely and kept warm until they regained mobility. Once stable, the mice were transferred to the telemetry room of the Animal Research Facility, where continuous data were collected via a dedicated computer running Dataquest software. The mice were allowed a 5–7 day recovery period to equilibrate the telemetry system and ensure accurate cardiovascular phenotyping. The mice had free access to food and water and housed in a temperature-controlled facility with a 12-hour light/dark cycle, and ambient temperature kept between 20°C and 26°C.

### Immunoblotting

Flash-frozen kidney tissue samples were homogenized in RIPA buffer using a tissue homogenizer (Thermo Fisher Scientific). The homogenized samples were centrifuged at 1300 rpm for 15 minutes at 4°C, and the supernatants were carefully collected. The protein concentrations in the supernatants were measured using the Bradford assay (Sigma-Aldrich) with results read on a Synergy Mx plate reader (BioTek) at 595 nm. Equal amounts of protein (40 µg per sample) were denatured by adding Laemmli buffer (Sigma-Aldrich) and heated at 95°C for 5 minutes. The denatured proteins were resolved by electrophoresis on 12% Criterion polyacrylamide gels (Bio-Rad) under reducing conditions. Following electrophoresis, the proteins were transferred onto nitrocellulose membranes (Bio-Rad) for electroblotting.

The membranes were blocked by incubation with 5% non-fat milk in 1x Tris-buffered saline (TBS) with 0.05% Tween-20 (TBS-T) for 1 hour at room temperature, with gentle shaking at 100 rpm. The membranes were then incubated overnight at 4°C with primary antibodies diluted in blocking buffer. The following primary antibodies were used: GRK4 (≈ 60 kDa, Sc-37689, mouse monoclonal, 1:250, Santa Cruz Biotechnology), SLC4A5 (≈ 150 kDa, PA5-113404, rabbit polyclonal, 1:500, Invitrogen), NHE3 (≈ 75 kDa, SC-136368, Mouse monoclonal, 1:500, Santa Cruz Biotechnology), HDAC1 (≈ 65 kDa, PA1-860, rabbit polyclonal, 1:1000, Invitrogen). GAPDH (≈ 37 kDa, MAB374, mouse monoclonal, 1:1000, Sigma Aldrich) as housekeeping proteins. The relative expression of target proteins was normalized to GAPDH as the loading control. The band intensities were quantified using ImageJ software (National Institutes of Health) and expressed as fold-change relative to control samples.

#### Quantitative RT-PCR

RNA was extracted from mouse renal cortex using the RNeasy Kit (Qiagen, #74104) as per the manufacturer’s guidelines. The RNA was quantified using a NanoDrop 2000 spectrophotometer (Thermo Fisher). cDNA was synthesized with the iScript™ cDNA Synthesis Kit (BioRad, #1708891), and quantitative PCR was carried out on an iQ™ 5 Real-Time PCR System (BioRad) with SYBR Green mix (Takara). Prevalidated KiCqStart primers (Sigma-Aldrich) were used to amplify mouse genes of interest, including *GRK4* (NM_019497), *NHE3* (NM_001081060), *NHE1* (NM_016981), *SLC4A5* (NM_001166067), *Na/K ATPase* (NM_144900), *SLC12A1* (NM_183354), *SGLT2* (NM_133254), and 𝛼*-ENaC* (NM_011324).

β-actin (NM_007393) served as the reference gene. All primers were purchased from Qiagen. The reactions utilized RT² SYBR Green ROX qPCR Mastermix (Qiagen), and the data were analyzed via the ΔΔCt method.

### Immunohistochemistry

The expression of GRK4 in mouse kidneys was analyzed using immunohistochemistry. The kidneys were harvested and fixed in 3.7% paraformaldehyde for 1 hour at 4°C. After fixation, the tissues were washed in phosphate-buffered saline (PBS) three times for 5 minutes each and then equilibrated overnight in 30% sucrose in PBS. Six-micrometer samples were prepared, and the slides were deparaffinized in xylene at 60°C for 1 hour and then washed twice for 3 minutes in 100%, 95%, and 75% ethanol. The slides were boiled in sodium citrate buffer (pH 6.0) for 3 minutes under pressure for antigen retrieval, cooled to room temperature, and then blocked in 1% BSA for 30 minutes at room temperature.

The sections were immunostained for GRK4 (Sc-37689, mouse monoclonal, 1:250, Santa Cruz Biotechnology) for 24 hours at 4°C using Alexa fluor® 647 donkey vs. mouse secondary antibody. Anti-SLC4A5 rabbit polyclonal antibody (PA5-87897, rabbit polyclonal, 1:200, Invitrogen) followed by Alexa Fluor® 647 donkey vs. rabbit secondary antibody. DAPI was used to visualize the nuclei. Confocal images were obtained sequentially in separate channels to avoid bleed-through using a Carl Zeiss LSM 510 META with a ×63/1.4 NA oil-immersion objective (Carl Zeiss, Thornwood, NY, USA) and processed using Zeiss 510 META with Physiology 3.5 and Multiple Time Series 3.5 software.

### Cell models

#### Preparation of Human Renal Proximal Tubule Cells (hRPTCs) with GRK4 Variants

Kidney tissues were collected in accordance with protocols approved by the University of Virginia Institutional Review Board and in compliance with the ethical principles outlined in the Declaration of Helsinki. hRPTCs were isolated from kidney specimens of patients who had undergone unilateral nephrectomy due to renal carcinoma or trauma. Only the visually and histologically normal sections of the kidney, located distal to the tumor or damaged area, were selected to ensure the use of healthy tissue for cell isolation (53, 63, 64).

The primary cultures of hRPTCs were subsequently immortalized to create stable cell lines, following established protocols (53, 63, 64). After immortalization, the hRPTCs were genotyped to identify specific variants in the GRK4 R65L (rs2960306), A142V (rs1024323), and A486V (rs1801058) and SLC4A5 (rs10177833 and rs7571842) genes. Based on the genotyping results, four distinct hRPTC lines were established, each carrying a unique combination of *GRK4* and *SLC4A5* variants: WT (*hGRK4γ WT* and *SLC4A5 WT*), SLC4A5 variant (*hGRK4γ WT* and *SLC4A5* variants), GRK4γ 65L (*hGRK4γ 65L* and *SLC4A5 WT*), and SLC4A5 GRK4γ 65L (*hGRK4γ 65L* and *SLC4A5* variants).

The cells were cultured under nonpolarizing conditions in a medium consisting of DMEM-F12 (Gibco), plasmocin (2.5 μg/ml, Fisher Scientific), epidermal growth factor (EGF, 10 ng/ml, Sigma), dexamethasone (36 ng/ml, Sigma), triiodothyronine (2 ng/ml, Sigma), insulin-transferrin-selenium (ITS, 1X, Invitrogen, 5 mL), penicillin/streptomycin (1X, Invitrogen, 10 mL), fetal bovine serum (2%, Invitrogen, 10 mL), and G418 (0.4%, EMD Chemicals).

### Cellular procedures

#### Quantitative RT-PCR

RNA was isolated from cells using the RNeasy Kit (Qiagen, #74104) and quantified with a NanoDrop 2000 spectrophotometer (Thermo Fisher), as above.

#### Immunoblotting

The cell samples were homogenized using a tissue homogenizer in RIPA buffer (Thermo Fisher Scientific) to ensure a homogenous mixture. The homogenized samples were prepared for immunoblotting, as described above.

#### Transwell Na^+^ accumulation assay

Intracellular free sodium ([Na^+^]i) was measured in hRPTCs using the cell-permeant dye Sodium Green™ tetraacetate (Molecular Probes, Eugene, OR, USA) (63). The hRPTCs were cultured to confluence under polarized conditions on polyester membrane Transwell-Clear inserts (Corning, Lowell, MA, USA) in 12-well plates. Transepithelial electrical resistance was monitored using an Epithelial Voltohmmeter to ensure confluency.

The hRPTCs were treated on the apical side and basolateral side with vehicle or 50 μM ouabain (CAS: 11018-89-6, PHR1945, Sigma Aldrich) at the basolateral side for 60 minutes to inhibit Na^+^/K^+^-ATPase, 100 nM 5-(N-ethyl-N-isopropyl)amiloride (EIPA) (CAS: 1154-25-2, A3085, Sigma Aldrich) at the apical side for 60 minutes to inhibit NHE3 activity, and 1 μM fenoldopam mesylate (CAS: 67227-57-0, 1269458, Sigma Aldrich) at the apical or basolateral side for 15 minutes. Sodium Green™ was then added to both sides for 30 minutes. Fluorescence emission was measured using a Victor II plate reader (excitation at 485 nm and emission at 535 nm). Protein concentrations were measured with the Pierce BCA Protein Assay Kit (Thermo Fisher Scientific, Rockford, IL, USA) and used to normalize the results. Results are reported as fluorescence units (Victor II reading) per mg of protein, and the percentage of inhibition or activation was calculated.

#### Statistics

All data are presented as mean ± standard error of the mean (SEM). Normality was determined with D’Angostino and Pearson omnibus K2 normality test or Saphiro-Wilk normality test, followed by the corresponding parametric or non-parametric test. P value < 0.05 was considered statistically significant. Statistical analyses were performed using GraphPad Prism 9.5.1 for Windows (GraphPad Software, San Diego, California USA).

#### Animal Study approval

All animal protocols used in this study were conducted in accordance with the NIH guidelines and were approved by the Institutional Animal Care and Use Committee at the George Wasdhington University.

## 3. Results

### 3.1 Human variant *GRK4 65L* induces salt sensitivity in mice

We have reported that a high-salt diet increases BP and renal GRK4 protein expression in salt-sensitive C57Bl/6J mice but not in salt-resistant SJL/J mice (35). The salt sensitivity of C57Bl/6J mice, which is associated with increased renal expression of GRK4, is replicated in our current report (**Fig. 2A**). In mice on C57Bl/6J background, the global, germline deletion of *Grk4* (*mGrk4^-/-^*) decreased systolic and diastolic BPs, in agreement with our previous report (27) and became salt resistant (Fig. 2A).

**Figure 1.**
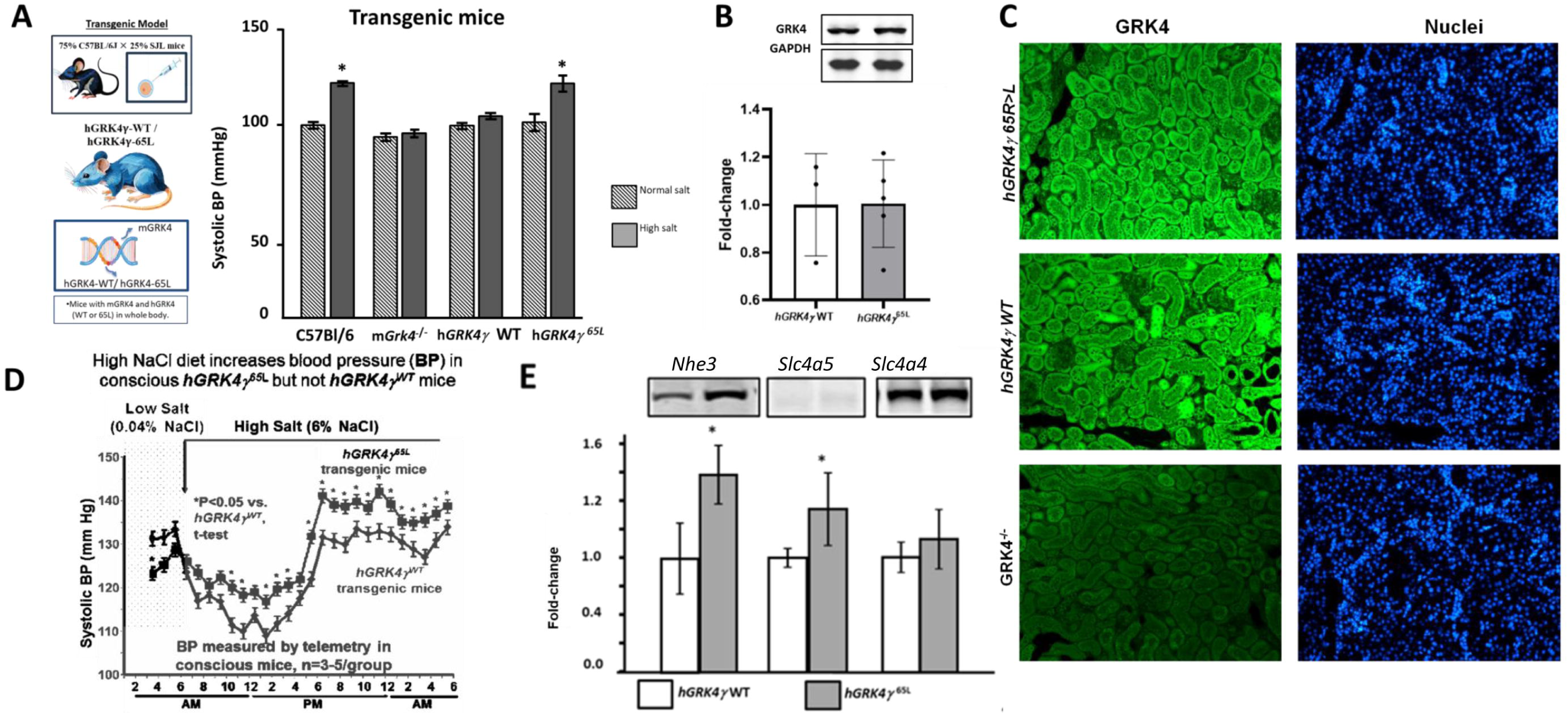
The human GRK4 65L variant induces salt sensitivity in mice. Blood Pressure and Renal GRK4 and Sodium Transporter Expressions in hGRK4 Transgenic Mice under Varying Salt Diets: **A)** Systolic BP (mmHg) measured from the femoral artery under anesthesia in 75-25%C57Bl/6J/SJL, mGrk4^-/-^, hGRK4γ WT and hGRK4γ-65L mice, *P<0.05, n=3-6/group, one-way ANOVA, Holm-Sidak test. *(B)* Telemetry-based BP monitoring in conscious hGRK4γ WT and hGRK4γ 65L mice after 7 days of low-salt diet (0.04% NaCl) and transitioned to a high-salt diet (6% NaCl); each point represents a 7-day average (\**P*<0.05, n=3-5/group, one-way ANOVA, Holm-Sidak test). *(C)* Renal GRK4 expression in hGRK4γ WT and hGRK4γ 65L transgenic mice, (n=3-4/group). *(D)* Immunohistochemistry of GRK4 in hGRK4γ WT, hGRK4γ 65L, and mGRK4^-/-^ mice, visualized with a GRK4 mouse monoclonal antibody (1:250) and Alexa Fluor 488 secondary antibody; nuclei stained with DAPI (Scale bar = 10 μm). *(E)* Western blot analysis of renal Na^+^/H^+^ exchanger 3 (NHE3), *SLC4A5*, and *SLC4A4* proteins in hGRK4γ WT and hGRK4γ 65L mice normalized with GAPDH as the loading control. (\**P*<0.05, n=4/group, Student’s t-test)

**Figure 2.**
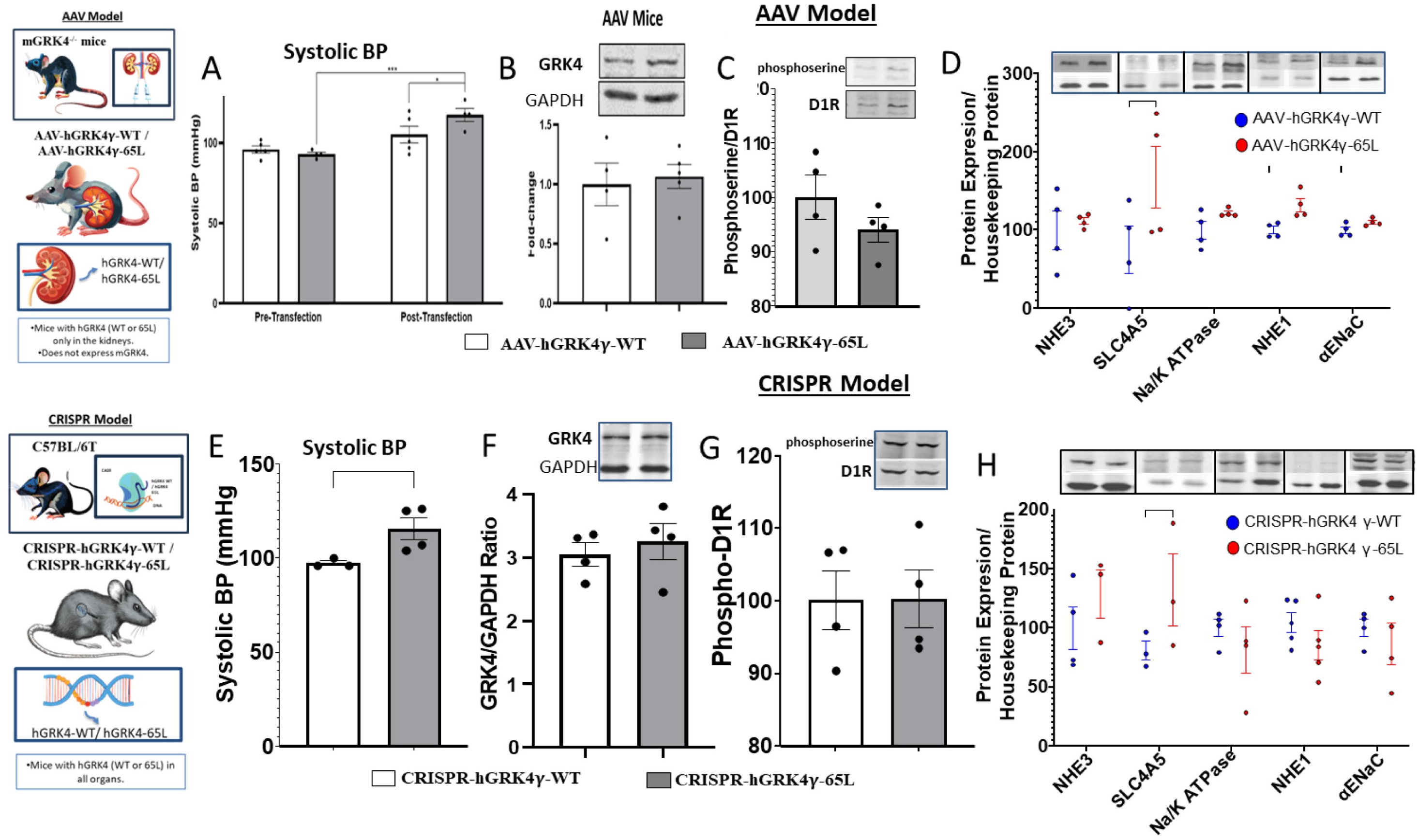
Comparison of Blood Pressure, Renal GRK4 Expression, and Renal Sodium Transporters/Pump in CRISPR and AAV Models for Evaluating GRK4 Variants: *(A)* Systolic blood pressure (BP), measured via the carotid artery, in pentobarbital-anesthetized GRK4 knockout (GRK4 KO) mice before and after bilateral ureteral transfection with AAV vectors carrying either hGRK4γ WT (AAV-hGRK4γ WT) or hGRK4γ 65L (AAV-hGRK4γ-65L), (*P*<0.05, n=3/group, one-way ANOVA, Holm-Sidak test *(B)* Western blot analysis of GRK4 protein expression, normalized by GAPDH as the loading control, in AAV-transfected mice shows no difference between AAV-hGRK4γ WT and AAV-hGRK4γ 65L mice (*P*<0.05, n=4/group, Student’s t-test). *(C)* D1 receptor (D1R) activity, represented by the phosphoserine D1R and D1R ratio, shows no significant difference between AAV-hGRK4γ WT and AAV-hGRK4γ-65L mice (*P*>0.05, n=4/group, Student’s t-test). *(D)* Western blots a of NHE3, SLC4A5, Na^+^/K^+^ ATPase, NHE1, and α-ENaC in AAV-hGRK4γ WT and AAV-hGRK4γ 65L mice normalized with GAPDH as the loading control (*P*<0.05, n=4/group, Student’s t-test). *(E)* Systolic BP in CRISPR-generated hGRK4γ WT and hGRK4 65L mice, shows a significant BP increase in mice with the 65L variant (*P*<0.05, n=3-4/group, Student’s t-test). *(F)* Western blots of GRK4 protein in CRISPR-hGRK4γ-WT and CRISPR-hGRK4γ-65L mice, (*P*<0.05, n=4/group). *(G)* D1R activity, represented by the phosphoserine/D1R ratio is not different between CRISPR-hGRK4γ WT and CRISPR-hGRK4γ 65L mice (*P*<0.05, n=4/group, Student’s t-test). *(H)* Western blots of NHE3, SLC4A5, Na^+^/K^+^ ATPase, NHE1, and α-ENaC in CRISPR-hGRK4γ WT and CRISPR-hGRK4γ 65L mice normalized with GAPDH as the loading control (*P*<0.05, n=4/group, Student’s t-test).

As aforementioned, in humans, the three GRK4γ variants, GRK4γ 65L, GRK4γ 142V, and GRK4γ 486V have been reported to be associated with hypertension with or without salt sensitivity, in several ethnic groups (14, 19, 24, 25).To elucidate the mechanisms underlying the GRK4 65L-mediated salt sensitivity and hypertension, we studied mice in which the human *GRK4*𝛾 *WT* (hGRK4γ WT) or the human *GRK4*𝛾 *65L* (rs2960306) transgenes were expressed under a CMV promoter while the mouse GRK4 was also expressed (25,26). Similar to the *Grk4^-/-^* mice, transgenic *hGRK4*𝛾 *WT* mice were salt-resistant whereas *hGRK4*𝛾 *65L* mice were salt-sensitive (**Fig. 1A**). Renal GRK4 protein levels were comparable between hGRK4γ WT and hGRK4γ 65L mice (**Fig. 1B**). Immunofluorescence of renal GRK4 expression and distribution were similar in hGRK4γ-WT and hGRK4γ-65L mice, while mGRK4^-/-^ mice had minimal GRK4 fluorescence (**Fig. 1C**). The circadian rhythm of BP was not affected in these mice (**Fig. 1D**). However, the renal expressions of NHE3 (sodium hydrogen exchanger type 3) and SLC4A5 (electrogenic sodium bicarbonate cotransporter 4) but not SLC4A4 (electrogenic sodium bicarbonate cotransporter 1) were higher in hGRK4γ 65L than hGRK4γ WT mice (**Fig. 2E**). (**Fig. 1E**)

To investigate further the difference in salt sensitivity between transgenic hGRK4γ WT and hGRK4γ 65L mice, we generated two other mouse models. The first model was generated by the retrograde bilateral ureteral infusion of adeno-associated (AAV)-9 vectors carrying either *hGRK4*𝛾 *WT* (AAV-*hGRK4*𝛾 *WT*) or *hGRK4*𝛾 *65L* gene (AAV-*hGRK4*𝛾 *65L*) in mGrk4^-/-^ mice. This model allowed us to compare the data from transgenic mice that express the *hGRK4γ WT* or *hGRK4*𝛾 *65L* only in the kidney with the rest of the body not expressing *mGrk4*. The second mouse model was generated using CRISPR/Cas9 technology to knock-in the human *GRK4γ WT* or *GRK4γ 65L* gene in the entire body while deleting the endogenous m*Grk4* gene in the entire body.

The BP before the retrograde ureteral AAV-mediated transfection was similar in the two groups of mGRK4^-/-^ mice fed a normal salt diet. However, 14 days after AAV-mediated transfection, the BP increased in AAV-hGRK4γ 65L but not in AAV-hGRK4γ WT mice (**Fig. 2A**). Thus, the expression of hGRK4γ 65L, only in the kidney, can increase BP even when sodium intake is not increased.

The higher BP in AAV-hGRK4γ 65L mice fed a normal salt diet was not related to increased renal expression of GRK4 because GRK4 expression was similar in the kidney cortex of mice transduced with AAV-hGRK4γ WT and AAV-hGRK4γ 65L **(Fig. 2B**). AAV-hGRK4γ WT and AAV-hGRK4γ 65L mice also had similar levels of phosphorylated D_1_R but tended to be lower in the AAV-hGRK4γ 65L group (**Fig. 2C**). The renal expressions of NHE3, Na/K ATPase, NHE1, and αENaC were similar in both groups but SLC4A5 expression was higher in AAV-hGRK4γ 65L than in AAV-hGRK4γ WT mice fed a normal salt diet (**Fig. 2D**).

Similar to the AAV studies, the BPs were also higher in CRISPR-hGRK4γ-65L than CRISPR-hGRK4γ-WT (**Fig. 2E**). Also similar to the AAV mouse model, the renal expressions of GRK4 protein in CRISPR-hGRK4γ WT and CRISPR-hGRK4γ 65L were not different from each other (**Fig. 2F**). The expressions of phosphorylated D_1_Rs were also similar in CRISPR-hGRK4γ WT and CRISPR-hGRK4γ 65L mice (**Fig. 3G**). Also, similar to the AAV-hGRK4γ WT and AAV-hGRK4γ 65L mice, the renal expressions of NHE3, Na/K ATPase, NHE1, and αENaC were similar in both groups but SLC4A5 expression was higher in CRISPR-hGRK4γ 65L than CRISPR-hGRK4γ WT group (**Fig. 3H**).

**Figure 3.**
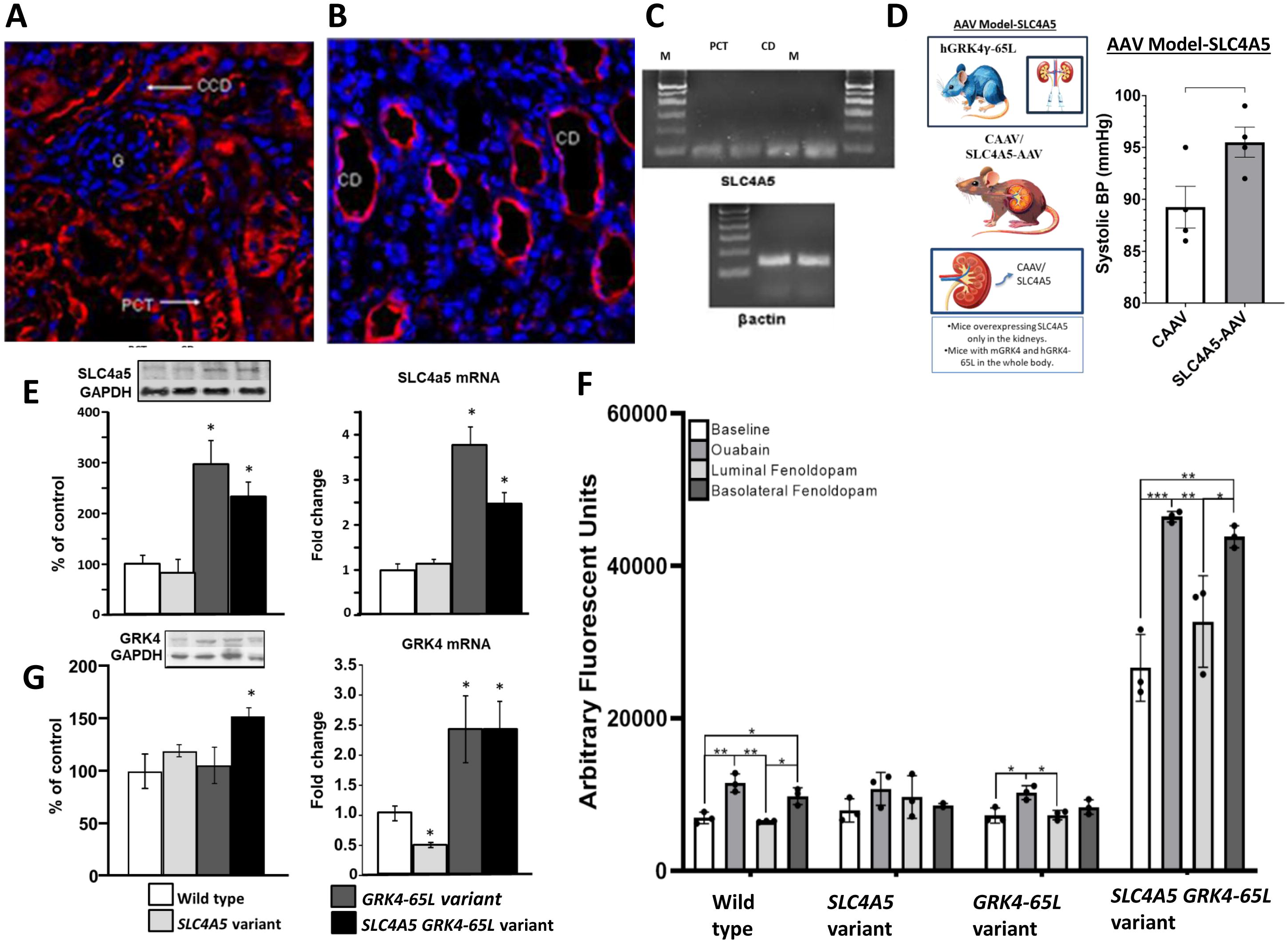
Analysis of SLC4A5 and GRK4 Expression and Function in Mouse and Human Renal Models. *(A)* Immunofluorescence staining for SLC4A5 (red) in the mouse renal cortex, showing expression in the proximal convoluted tubule (PCT), glomerulus (G), and cortical collecting duct (CCD).Blue = DAPI for nuclei (Scale bar = 10 μm). *(B)* Immunofluorescence staining for SLC4A5 (red) in the renal medulla, showing expression in the collecting duct (CD). *(C)* RT-PCR of *Slc4a5* mRNA expression in laser-captured microdissected PCT and CCD from mouse kidneys, shows mRNA in both tubule segments. *(D)* BPs (measured from the carotid artery of pentobarbital anesthetized mice) are higher in hGRK4γ 65L transgenic than CAAV mice fed a high salt (4 % NaCl) diet. Kidneys of the mice were transduced by the retrograde ureteral infusion of control AAV (CAAV) or AAV carrying human wild-type SLC4A5 (SLC:AAV-SLC4A5 1^11^ vgp /kidney) (*P<0.05 Student’s t-test, n=4/group). *(E)* Western blot and qPCR analyses of SLC4A5 protein and *Slc4a5* mRNA levels in human renal proximal tubule cells (hRPTCs) expressing wild-type (WT) SLC4A5, SLC4A5 variants (rs10177833 and rs7571842), GRK4 65L variant, and both SLC4A5 and GRK4 65L). GAPDH is used as the loading control (\**P*<0.05, n=3/group, Student’s t-test). *(F)* Western blots and qPCR analyses of GRK4 protein and *Grk4* mRNA levels in WT, SLC4A5 variants, GRK4-65L variant, and SLC4A5-GRK4-65L variants hRPTCs, with GAPDH as the loading control (*P*<0.05, n=3/group, Student’s t-test). *(G)* Na^+^ accumulation in hRPTCs: Bar graphs illustrate intracellular sodium (Na^+^) concentration (arbitrary fluorescence units) in WT, SLC4A5 variant, GRK4 65L variant, and SLC4A5-GRK4 65L hRPTCs treated with our without ouabain, the inhibitor of Na^+^/K^+^ ATPase and/or fenoldopam, D1R-like agonist. P<0.05, one-way ANOVA, Holm-Sidak test (*n*=3).

### 3.2 Increased SLC4A5 Activity is Associated with Salt Sensitivity in GRK4γ 65L mice

*SLC4A5 (NBCe2)* gene variants have been linked to increased blood pressure (BP) and/or salt sensitivity in several ethnic groups (24, 38–42). NBCe2 (39, 43) is present in a subapical compartment in human renal proximal tubule cells cultured in low NaCl concentration (120 mM) and migrates to the microvilli in high NaCl concentration (170 mM) (43). To investigate the underlying mechanisms of the increased SLC4A5 expression in *hGRK4*𝛾 *65L* mice, we studied the cellular expression of SLC4A5. Immunofluorescence microscopy showed the presence of SLC4A5 in the luminal membrane of the renal proximal convoluted tubule (PCT), cortical collecting duct (CCD) (**Fig. 3A**) and medullary collecting duct (CD) (**Fig. 3B**) in C57Bl/6J mice. Mass spectrometry confirmed the selectivity of the antibody to detect SLC4A5 protein (Supplemental Figure 1). RT-PCR of renal PCT and CD, obtained by laser capture, confirmed the expression of *Slc4a5* mRNA in the PCT and CCD (**Fig. 3C**), corroborating the immunostaining findings. This distinct renal tubule-specific distribution of *Slc4a5* suggests a tubule segment-specific regulation of the transport of sodium and bicarbonate along the nephron.

To assess the role of SLC4A5 on BP regulation, we developed a fourth animal model using transgenic hGRK4γ 65L mice with retrograde ureteral infusion of AAV vectors carrying either a control adeno-associated virus vector (CAAV) or the human wild-type *SLC4A5* gene (*SLC4A5*-AAV). Systolic BP was measured from the carotid artery under pentobarbital anesthesia in mice fed a high salt diet (4% NaCl) for 2 weeks. hGRK4γ 65L mice with kidneys transduced with *SLC4A5*-AAV had higher BP than those infused with CAAV (**Fig. 3D**). This demonstrated that an increase in renal SLC4A5 may, in part, be involved in the increase in BP in transgenic hGRK4γ 65L mice.

To explore further the effect of SLC4A5 and GRK4 65L on renal function, we studied human renal proximal tubule cells (hRPTCs) expressing either the SLC4A5 variants (rs10177833 and rs7571842), GRK4 65L variant, or both (39, 43). GRK4 protein expression was increased in hRPTCs expressing both *GRK4 65L* and *SLC4A5* variants (**Fig. 3E and 3G)**. However, *GRK4* mRNA was increased in the presence of *GRK4 65L* but not increased further by the presence of *SLC4A5* variants. By contrast, *GRK4* mRNA was decreased in hRPTCs expressing the *SLC4A5* variant. These results suggest that there is at some level an interaction between GRK4 and SLC4A5.

We then assessed the effects of the above genes on sodium transport in polarized hRPTCs grown in culture (**Fig. 3G**, **Table 2**). In all the cell types, treatment with ouabain increased intracellular sodium because of the inhibition of sodium transport at the basolateral side. However, the increase in intracellular sodium was least in the cells with SLC4A5 variant, suggesting increased activity of another sodium transporter in the basolateral membrane, e.g., NBCe1 (44). In wild-type cells, the D_1_ receptor agonist fenoldopam added on the luminal side reduced intracellular sodium, albeit minimally, relative to baseline (**Fig. 3F**, **Table 1**), likely through inhibition of NHE3 and SLC4A5 (**Fig. 3F**), while addition of fenoldopam on the basolateral side increased sodium accumulation, likely through inhibition of Na^+^/K^+^ ATPase activity (**Fig. 3F**). Treatment of the hRPTCs expressing either the SLC4A5 variant or GRK4 65L with fenoldopam at the basolateral or luminal side had no effect of intracellular sodium indicating that sodium transporters are not responsive to D_1_-like receptor inhibition in the presence of these variants.

Notably, sodium transport was markedly altered in hRPTCs co-expressing both variants. Baseline sodium accumulation was three times higher in hRPTCs co-expressing both variants than in the other cell lines. Treatment with ouabain alone markedly increased intracellular sodium, indicating increased luminal sodium transport. The treatment with fenoldopam on the basolateral side increased sodium accumulation to a greater extent in the presence of both SLC4A5 and GRK4γ 65L than in the absence or presence of only *SLC4A5* variants or *GRK4* variants, indicating increased sodium transport at the luminal side in addition to the minimal inhibition of sodium transport by fenoldopam at the basolateral side. Treatment with fenoldopam on the luminal side also decreased intracellular sodium but not below the baseline level as seen in the hRPTCs without gene variants, indicating impaired inhibitory effect of fenoldopam at the luminal side. These findings suggest that the combination of the SLC4A5 and GRK4γ 65L variants contributes to hypertension by increasing basolateral and luminal sodium transport and impairing the inhibition of sodium transport mediated by D1-like receptor agonist, fenoldopam at the luminal and basolateral sides. This disruption of sodium homeostasis may have contributed to the observed elevation in BP in the combined presence of SLC4A5 and GRK4γ 65L variants.

### 3.3 HDAC1 expression is decreased in GRK4 65L mice and hRPTCs with both GRK4 65L and SLC4A5 variants but not in hRPTCs with GRK4 65L, alone

We have reported that renal GRK4 can negatively regulate the expression and function of HDAC1 in the kidney (27). HDACs cut out the acetyl groups in histones increasing their binding to DNA, resulting in a decreased gene transcription (45). In hRPTCs, the expression of an activating variant of GRK4, i.e., GRK4 142V, increased HDAC1 phosphorylation, decreased HDAC1 nuclear activity, and increased the transcription of AT_1_R. Furthermore, in mice, renal-selective silencing of HDAC1 increased BP (27). We found that renal HDAC1 mRNA is decreased in salt-sensitive transgenic hGRK4γ 65L mice fed a high-salt diet (**Fig 4A**). Similarly, mice with AAV-transduced renal expression of hGRK4γ 65L (**Fig 4B**) or CRISPR-hGRK4γ 65L mice (**Fig 4C**) also on high salt diets, have decreased renal HDAC1 protein expression. Moreover, HDAC1 phosphorylation in cells expressing the GRK4γ 65L (and maybe SLC4A5) alone or both GRK4γ 65L and SLC4A5 was increased, indicating decreased nuclear activity (**Fig 4D**). These effects are specific to HDAC1 because the phosphorylation of HDAC2 was similar in all groups (**Fig 4E**). By contrast in hRPTCs, HDAC1 protein expression was not different between those carrying GRK4 65L and GRK4 WT but lower in hRPTCs expressing the SLC4A5 variant alone or in combination with GRK4γ 65L (**Fig 4H**), suggesting that both these variants are associated with decreased HDAC1 expression that is more prominent in humans than in mice (**Fig. 4D**).

**Figure 4.**
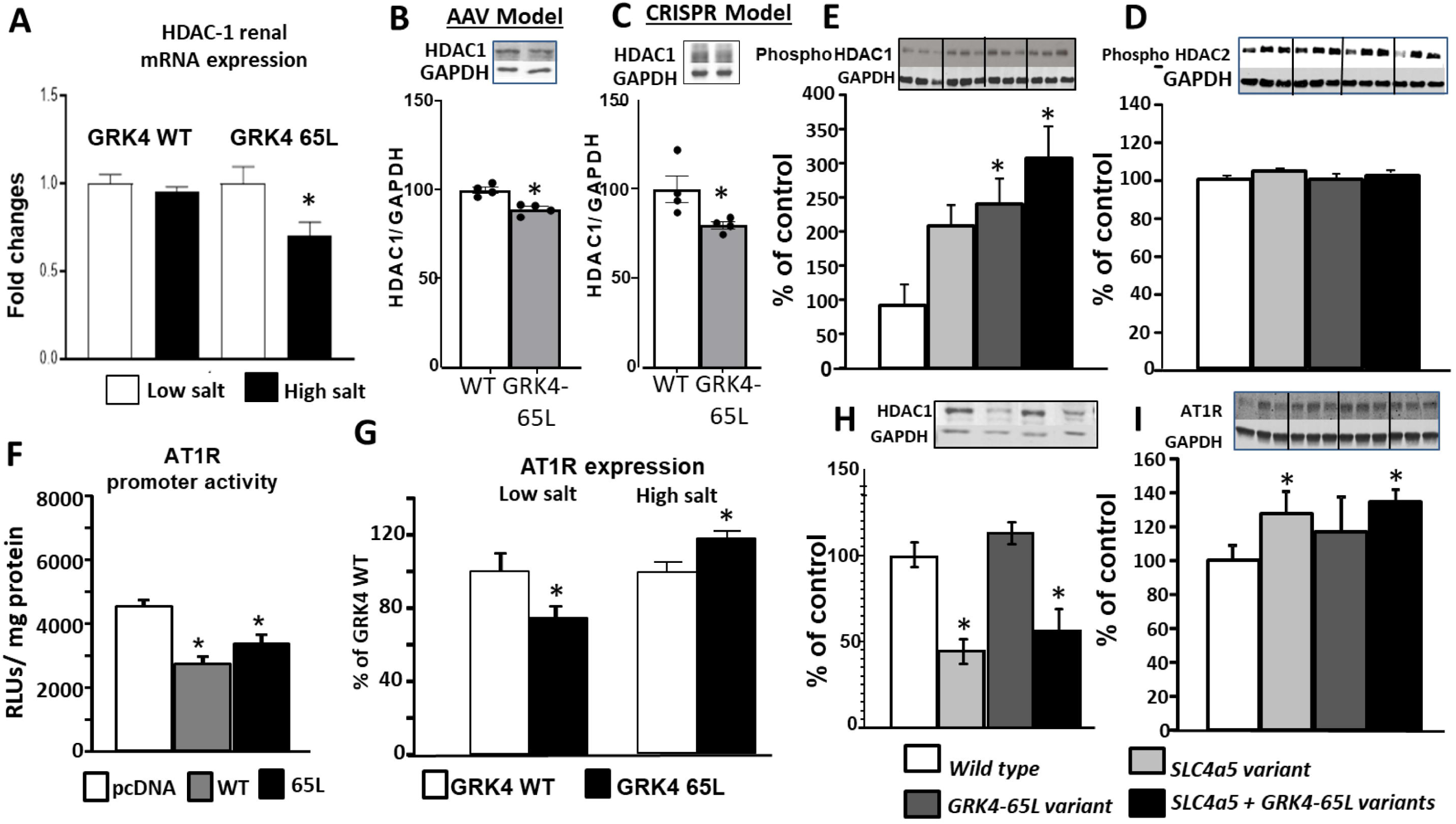
Mechanistic Role of GRK4 WT and GRK4 65L Variant in Regulating HDAC1 Expression and Sodium Transport in Renal Models: (A) qRT-PCR of renal HDAC1 mRNA in hGRK4γ WT and hGRK4γ 65L transgenic mice on normal-and high-salt diets (*P*<0.05, *n*=3/group, Student’s t-test). (B) Western blots of renal HDAC1 protein in AAV-hGRK4γ WT and AAV-hGRK4γ 65L mice fed a high salt (6% NaCl) diet for one week, (GAPDH is the loading control; *P*<0.05, *n*=4/group, Student’s t-test). (C) Western blots of renal HDAC1 in CRISPR-hGRK4γ WT and CRISPR-hGRK4γ 65L mice also fed a high salt diet for 7 days (GAPDH is the loading control; *P*<0.05, *n*=4/group, Student’s t-test). (D) Ratio of phosphorylated to total HDAC1 protein in hRPTCs with WT SLC4A5 and GRK4, SLC4A5 variant, GRK4 65L, and SLC4A5 variant-GRK4 65L groups (GAPDH is the loading control; *P*<0.05, *n*=3-5/group,ANOVA). (E) Ratio of phosphorylated to total HDAC2 protein in hRPTCs, analyzed as above (GAPDH is the loading control; *P*<0.05, *n*=3-5/group, ANOVA. (F) Luciferase assay 24h after transfection of hRPTCs with pAT_1_RLuc plasmid (human AT_1_R promoter, from +281 to-1941 bp, driving the expression of firefly Luciferase) and pcDNA, hGRK4𝛾 WT or hGRK4𝛾 65L (*P<0.01 vs. pcDNA, n=4/group; one-way ANOVA, Holm-Sidak test). (G) Renal AT_1_R mRNA expression in hGRK4 65L and GRK4𝛾 WT transgenic mice on low (0.09%) or high (4%) NaCl diet (P<0.05 vs. hGRK4WT, n=3/group, Student’s t-test). (H) Western blots of HDAC1 in hRPTCs expressing *GRK4 WT* and *SLC4A5 WT*, *SLC4A5* variant, *GRK4 65L* variant, and both *SLC4A5* and *GRK4 65L* variants, (GAPDH is the loading control; *P*<0.05, *n*=3-5/group, ANOVA). (i) Western blots of AT_1_R protein in hRPTCs from *GRK4 WT* and *SLC4A5 WT, SLC4A5* variant, *GRK4-65L* variant, and both SLC4A5 GRK4-65L variants. GAPDH was used as a loading control (P<0.05, n=3/group, one-way ANOVA, Holm-Sidak test).

The increased renal expression of AT_1_R protein in GRK4γ 65L mice on high salt diet and hRPTCs bearing GRK4 65L variant may be related to a decrease in HDAC1 (**Figs. 4B** and **4C**) that is also found in hRPTCs with both GRK4γ 65L and the SLC4A5 (**Figs. 4H** and **4I)**. To assess further the role of HDAC1 in increasing AT_1_R expression in these cells, we studied the AT_1_R promoter activity in hRPTCs transfected with GRK4γ WT or GRK4γ 65L and found that in contrast with what we reported for the GRK4γ 142V (26), the GRK4γ 65L variant did not increase but instead decreased the activity of the AT_1_R promoter (**Fig 4F**). However, the transcription of the AT_1_R in transgenic mice expressing hGRK4γ WT or hGRK4γ 65L was dependent on the salt intake. On a low salt diet AT_1_R mRNA was lower in hGRK4γ 65L than in hGRK4γ WT mice but on a high salt diet AT_1_R mRNA was higher in hGRK4γ 65L than hGRK4γ WT mice (**Fig. 4G**). These results support the role of HDAC1 in the salt sensitivity of these mice that may be related to the increase in AT_1_R expression.

## 4. Discussion

This report analyzed for the first time the phenotypic consequences of the human GRK4γ 65L variant and investigated the molecular mechanisms involved in the salt-sensitive hypertension associated with the presence of this variant. Our preliminary studies showed that the expression of hGRK4 65L in the entire body or only in the kidneys of mice causes salt-sensitive hypertension (46). Our new data show that GRK4γ 65L inhibits HDAC1 expression and activity, which should increase DNA acetylation and promote the expression of genes such as SLC4A5 and AGTR1. Electrogenic sodium bicarbonate cotransporter 4 (SLC4A5) increases the transport of sodium in the apical membrane of renal proximal tubule cells, reducing the ability to excrete sodium and subsequently increasing BP (39, 43).

GRK4, unlike the other six members of the GRK family, is mainly expressed in the testes and myometrium and to lesser extent in the brain, intestine, and kidney (13–20). GRK4 expression and activity in the kidneys and arteries is increased in the hypertensive state, relative to the normotensive state (19, 36–38). Several SNPs in the GRK4 gene, R65L, A142V, and A486V, have been associated with conditions such as hypertension and other adverse cardiovascular outcomes in humans (39), rodents (19, 23, 26–28, 54), and even zebrafish (13). To confirm the blood pressure phenotype of the GRK4 65L variant, we studied 3 different mouse models expressing the human GRK4γ 65L variant in the kidney. All the animal models developed salt-sensitive hypertension with normal BP when salt intake was normal. Similarly, previous studies have shown that the GRK4 65L genotype is inversely associated with hypertension risk and high salt diet in a cohort of 1394 Korean adults selected from a community-based cohort of 15,034 patients (40). Interestingly, the GRK4 65L allele has been reported to be associated with stress-induced reduction in urinary sodium in black normotensive adolescents (41). The GRK4 A486V polymorphism is also associated with obesity induced by high sodium intake n normotensive Korean children, but in this study the GRK4 A486V homozygous mutants were observed in boys and the GRK4 A486V heterozygote mutants were observed only in girls (42), suggesting that the effects of GRK4 SNPs on BP regulation may also depend on other factors such as age, sex, and race. In this regard, other studies have shown the effects of lifestyle modification on human GRK4 variants on BP response, showing that a low-salt diet in patients carrying GRK4 R65L showed a significant reduction in ambulatory BP (50), consistent with the data we found in GRK4g 65L transgenic mice (36). In this respect, other studies have reported the association of *GRK4* variants in the antihypertensive response to β-adrenergic receptor blockers. In both African-Americans and Caucasians, an increased number of copies of the GRK4 variant 65L-142V haplotype is linked to a decreased response to the β-blocker atenolol when used as monotherapy (39). Another study showed that hypertensive African-American men, but not women, with early hypertensive nephrosclerosis and GRK4 142V variant were less responsive to the β1-adrenergic receptor blocker metoprolol in the presence of the GRK4 65L variant (43)). Therefore, our mice data are consistent with the human data showing that GRK4 65L variant causes hypertension only when sodium intake is increased, similar to what has been reported for GRK4γ A486V transgenic mice (28). By contrast, the GRK4 A142V variant can cause hypertension even when sodium intake is normal (10, 25). Thus, the presence or absence of hGRK4 variants may guide the therapy for hypertension (38), however, the mechanisms involved in the phenotypic differences of GRK4 variants are still unclear.

Previous studies have reported that targeted mutation of *Slc4a5* (NBCe2, NBC4) in mice induces arterial hypertension with compensated metabolic acidosis (44) *SLC4A5* variants (rs7571842 and rs10177833) and GRK4 65L are also strongly associated with salt sensitivity of BP in 2 separate Caucasian populations (23). Thus, SLC4A5 may be involved in the salt sensitivity associated with the presence of hGRK4γ 65L. Our results show that all animal models and human cells with hGRK4γ 65L variant have an increase in renal SLC4A5 expression (Figures 1, 2), which suggest an interaction between SLC4A5 with the GRK4 65L in the pathogenesis of hypertension.

In addition to the ability of GRKs to phosphorylate and regulate the activity of GPCRs, other actions of GRKs via the phosphorylation of other proteins involved in gene expression regulation have been described (11–14, 19, 56). Previous studies have shown that GRK2 regulates the early and late stages of cell migration in cancer through HDAC6 phosphorylation (45). GRK5 has been shown to be a nuclear HDAC5 kinase that causes cardiac hypertrophy independent of the function of GPCRs (46). Moreover, GRK4 inhibits autophagy and promotes apoptosis by increasing the phosphorylation of HDAC4 in cardiomyocytes (47). GRK4γ 142V promotes angiotensin II-mediated hypertension by inhibition of the renal HDAC1 (26). All these pieces of evidence show that GRK proteins may have additional function in gene regulation thought HDAC, independent of GPCR effects. Our results show that GRK4γ 65L does not affect the expression of HDAC1 but may decrease its activity (**Figs. 4D** and **4H**). However, the GRK4γ 65L-mediated decrease in HDAC1 activity was not associated with increase in AT_1_R promoter activity (Figure 5F), in contrast to what we found with GRK4γ A142V (27). Rather, the decrease in HDAC1 expression and activity was related to the presence of SLC4A5 variants (**Fig. 5H**). HDAC1 can regulate major endothelial functions such as angiogenesis, inflammatory signaling, redox homeostasis, and nitric oxide signaling (60), all associated with BP regulation (48). It is therefore proposed that the presence of the SLC4A5 variants results in a gain of function that favors the inhibition of HDAC1, which in turn induces a reduction in DNA methylation, thereby increasing *Agtr1*gene expression and interacting with GRK4 65L to increase renal sodium reabsorption and subsequently BP (Figure 5).

**Figure 5.**
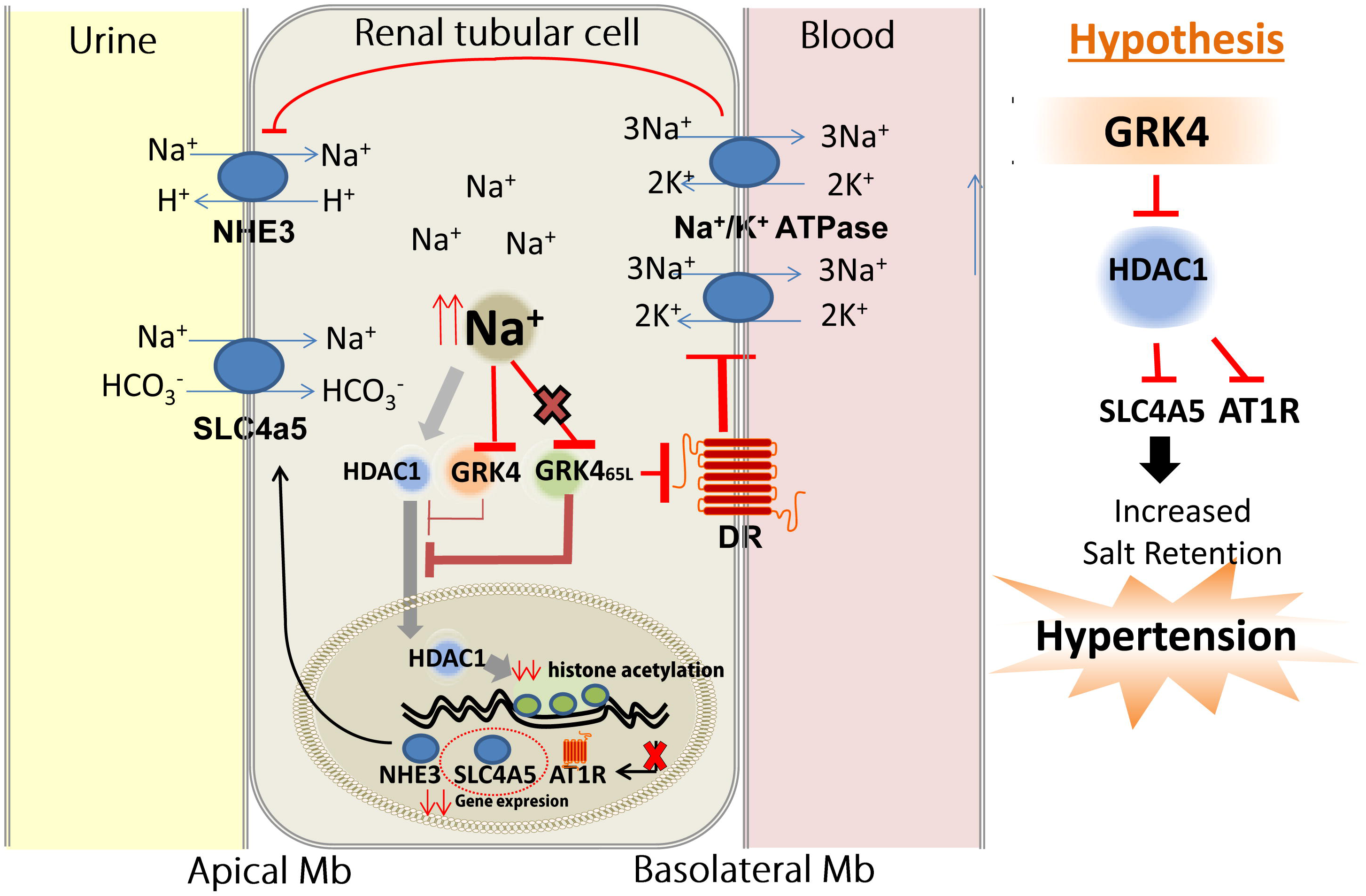
Graphical Abstract. This figure illustrates a renal proximal tubule cell depicting sodium (Na⁺) transport mechanisms through the apical and basolateral membranes and the hypothesized roles of HDAC1 (Histone Deacetylase 1) and GRK4 (G Protein-Coupled Receptor Kinase 4) in regulating sodium handling, potentially contributing to hypertension. The apical membrane, adjacent to the tubular fluid, features two key sodium transporters: NHE3 (Na⁺/H⁺ exchanger 3) and SLC4A5 (Solute Carrier Family 4 Member 5), which mediate Na⁺ reabsorption from the renal proximal tubular lumen. The basolateral membrane, interfacing with the blood, express Na⁺/K⁺-ATPase that pumps Na⁺ out of the cell into the bloodstream. It is hypothesized that an increase in intracellular Na⁺ levels activate two distinct pathways involving GRK4 and HDAC1. First, GRK4 activation negatively regulates dopamine receptor (DR) activity, impairing Na⁺/K⁺-ATPase function, leading to reduced Na⁺ transport out of the cell into the blood and further increasing intracellular Na⁺ levels. GRK4 variants, such as GRK4 65L, are proposed to fail in inhibiting HDAC1. Normally, HDAC1 activation removes the acetyl groups from DNA, which leads to tightening of chromatin, preventing the binding of transcription factors, suppressing the expression of genes, including SLC4A5, NHE3, and AT_1_R (Angiotensin II Type 1 Receptor). This epigenetic regulation reduces renal tubular Na⁺ reabsorption via SLC4A5 and NHE3 at the apical membrane, limiting Na⁺ uptake into the cell. However, the dysfunction of GRK4 in certain variants prevents this HDAC1 inhibition, increasing the expression of SLC4A5 and NHE3 and renal sodium reabsorption that is facilitated by AT1R, further compounding sodium imbalance. This coordinated dysregulation of signaling and sodium transport within the renal tubular cell is hypothesized to contribute to hypertension, with GRK4 playing a pivotal role in modulating both HDAC1 activity, Na⁺ transporters, and AT1R.

Our results highlight the underlying and compensatory renal mechanisms for the hypertension in mice with hGRK4γ 65L and the complex interaction between these variants in regulating not only SLC4A5 expression but also that of other genes involved in BP regulation such as AT_1_R and HDAC1. This new information clarifies the pathways involved in the salt sensitivity associated with hGRK4 gene variants, including GRK4γ 65L, which may have a significant effect in the prevention and treatment of hypertension.

## Data Sharing Statement

All data will be published following FAIR (Findable, Accessible, Interoperable and Reusable) principles. This will facilitate that the results of this study are easily reproducible, and that all the information generated (that does not affect sensitive data) can be easily found and reused in other studies.

## Supporting information

Datos suplementarios y leyendas de figuras

## Acknowledgments

The authors thank the Histology Services at the George Washington University Research Pathology Core Lab for their expertise.

## List of supplementary material

Including a Supplemental Figure S1. SLC4a5 is expressed in MRPTC and a supplemental Table S1. Effect of the D1-like receptor agonist, fenoldopam (1µM) on intracellular sodium concentration in human renal proximal tubule cells.

