## Supplementary material for "*The human GRK4γ 65L variant* causes salt-sensitive hypertension by increasing renal SLC4A5 expression through the HDAC1 pathway": Datos suplementarios y leyendas de figuras

**Supplementary Materials**

### GRK465L variant induces salt sensitivity hypertension via increases renal SLC4a5 expression thought HDAC1 pathway

Santiago Cuevas et al.

#### Supplemental Figures

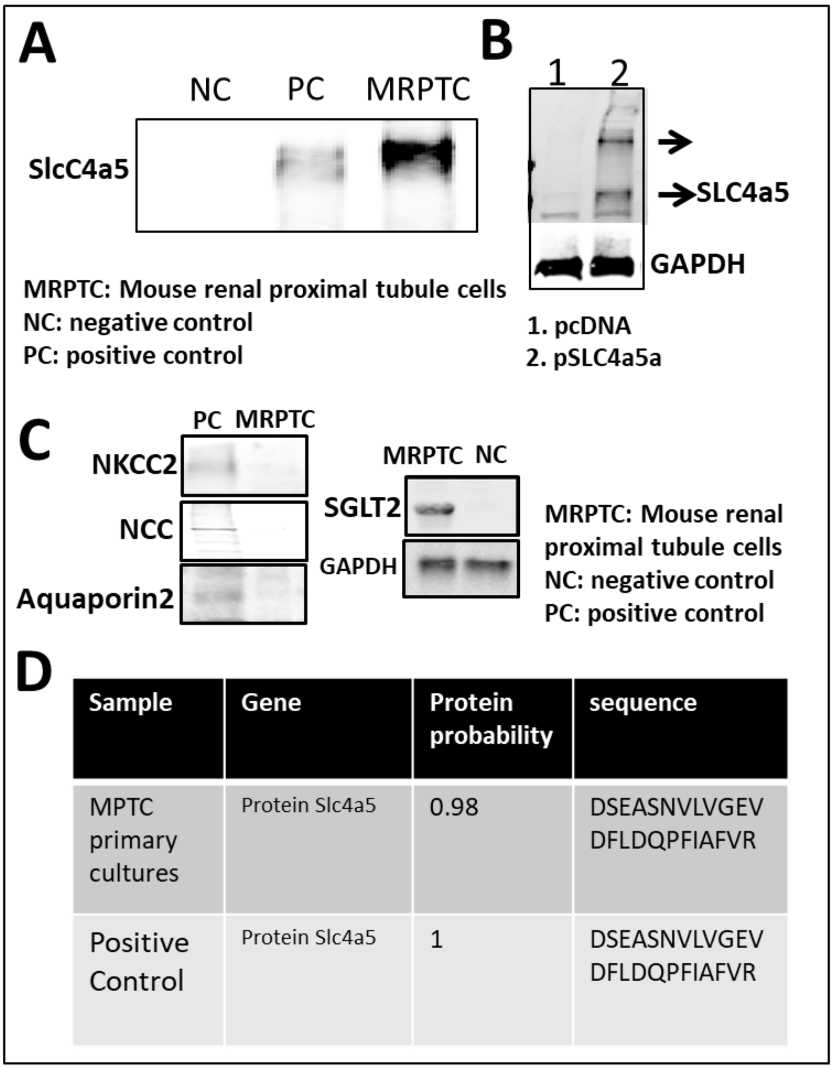


**Supplemental Figure 1 . SLC4a5 is expressed in MRPTC.** (**A**) Primary cultures of MRPTC (mouse renal proximal tubule cell) from BALB/c mice were immunoprecipitated with SLC4a5 (Sigma HPA036621) antibody and immunoblotted with SLC4a5 antibody. NC = negative control: primary cultures of MRPTC immunoprecipitated with rabbit IgG. PC= positive control: iNT16 (human renal proximal tubule cells) transfected with pSLC4a5 plasmid. (**B**) iNT16 cells transfected with pcDNA or pSLC4A5 plasmid. **(C)**. Homogenates of MRPTC primary cultures were immunoblotted using antibodies against NKCC2 (thick ascending limb), NCC (distal convoluted tubule), aquaporin 2 (principal cells of connecting tubule and collecting duct), and SGLT2 (S1 and S2 segments of proximal tubule). NC = negative control: primary cultures of collecting duct cells. PC= positive control: Total kidney homogenates. (**D**). Mass spectrometry analysis of the immunoblot bands corresponding to SLC4a5 from the lanes PC and MPTC (Figure A).

Table 1.

| Effect of the D1-like receptor agonist, fenoldopam (1 μM) on intracellular sodium concentration in human renal proximal tubule cells | | | | |
|---|---|---|---|---|
| | Wild-type | SLC4A5 variant | GRK4 65L | SLC4A5 variant + GRK4 65L |
| Baseline | 100±10 | 100±20 | 100±15 | 100±17 |
| Ouabain (50 μM) | 160±11 | 133±17 | 154±10 | 170±8 |
| Luminal fenoldopam | 86±4.5 | 115±15 | 100±3 | 119±30 |
| Basolateral fenoldopam | 136±10.5 | 100±5 | 115±5 | 162±10 |
